# Trophic Selective Pressures Organize the Composition of Endolithic Microbial Communities from Global Deserts

**DOI:** 10.1101/761262

**Authors:** Evan Qu, Chris Omelon, Aharon Oren, Victoria Meslier, Don A. Cowan, Gillian Maggs-Kölling, Jocelyne DiRuggiero

## Abstract

Studies of microbial biogeography are often convoluted by extremely high diversity and differences in microenvironmental factors such as pH and nutrient availability. Desert endolithic (inside rock) communities are exceptionally simple ecosystems that can serve as a tractable model for investigating long-range biogeographic effects on microbial communities. We conducted a comprehensive survey of endolithic sandstones using high-throughput marker gene sequencing to characterize global patterns of diversity in endolithic microbial communities. We also tested a range of abiotic variables in order to investigate the factors that drive community assembly at various trophic levels. Macroclimate was found to be the primary driver of endolithic community composition, with the most striking difference witnessed between hot and polar deserts. This difference was largely attributable to the specialization of prokaryotic and eukaryotic primary producers to different climate conditions. On a regional scale, microclimate and properties of the rock substrate were found to influence community assembly, although to a lesser degree than global hot versus polar conditions. We found new evidence that the factors driving endolithic community assembly differ between trophic levels. While phototrophic taxa were rigorously selected for among different sites, heterotrophic taxa were more cosmopolitan, suggesting that stochasticity plays a larger role in heterotroph assembly. This study is the first to uncover the global drivers of desert endolithic diversity using high-throughput sequencing. We demonstrate that phototrophs and heterotrophs in the endolithic community assemble under different stochastic and deterministic influences, emphasizing the need for studies of microorganisms in context of their functional niche in the community.

## Introduction

Arid and hyper-arid regions of Earth pose a significant challenge to life, with stresses such as high solar radiation, sparse and intermittent rain events, and drastic temperature fluctuations exerting strong selective pressures on organisms (Lebre et al., 2017; Potts and Webb, 1994). As aridity increases in an area and the diversity of higher eukaryotic life decreases, microorganisms begin to constitute a larger component of the total biomass and provide essential ecosystem functions such as carbon fixation and biogeochemical cycling (Makhalanyane et al., 2015; Pointing and Belnap, 2012). Desert microorganisms colonize rock and soil matrix as an adaptation to the harsh environmental conditions, forming a category of microbial refuges that includes biological soil crusts, hypoliths (under rock), epiliths (above rock), and endoliths (within rock) (Pointing and Belnap, 2012).

Endolithic communities, which inhabit pore and fissure space within the interior of rocks, are perhaps the most xerotolerant of these microbial refuges, appearing in the driest environments on Earth such as the hyper-arid core of the Chilean Atacama Desert and the McMurdo Dry Valleys in Antarctica (Azua-Bustos et al., 2012; Friedmann and Ocampo, 1976). These communities usually colonize a thin layer underneath the surface of the rock and have been described in diverse substrates including sandstone (Friedmann et al., 1967; Walker and Pace, 2007), limestone (Wong et al., 2010), halite (de los Ríos et al., 2010), gypsum (DiRuggiero et al., 2013), ignimbrite (Wierzchos et al., 2013), and granite (de los Ríos et al., 2005). Depending on the nature of the substrate, the primary mode of community colonization can either be cryptoendolithic, inhabiting pores within the rock matrix or chasmoendolithic, inhabiting cracks and fissure space in the rocks (Golubic et al., 1981). Within these communities, a few core photoautotrophic taxa along with a variety of chemoheterotrophic taxa are found living in consortium. *Cyanobacteria* are typically the dominant phototroph, often belonging to the multi-resistant genus *Chroococcidiopsis* (Friedmann et al., 1967; Meslier et al., 2018; Wierzchos et al., 2018), although eukaryotic green algae have also been reported in sandstone communities (Walker and Pace, 2007), gypsum (Wierzchos et al., 2015) and halite communities from the Atacama Desert (Robinson et al., 2015). Diverse heterotrophic bacteria are found in the endolithic community, with the phyla *Actinobacteria, Proteobacteria, Chloroflexi, Bacteroidetes*, and *Deinococcus-Thermus* particularly represented (Meslier et al., 2018; Walker and Pace, 2007; Wierzchos et al., 2018).

Desert lithic habitats have been well-utilized for studies of microbial biogeography, as their low diversity, simple trophic structure, and similar microenvironment provides a tractable model when investigating spatial patterns in microbial community assembly (Caruso et al., 2011). These studies have found that environmental determinants, in particular climate, are the primary factors that organize microbial communities at a global and regional scale (Chan et al., 2012; Friedmann, 1980; Lacap-Bugler et al., 2017). Biotic interactions are also thought to influence community assembly, and endolithic *Cyanobacteria* have been reported to play a key role in determining overall community structure and function (Valverde et al., 2015). Conversely, it remains poorly understood how stochastic processes influence assembly within the lithic habitat. Early studies of endolithic sandstones found remarkably similar communities at global scales, which led to the hypothesis that these communities are seeded from a global metacommunity (Walker and Pace, 2007). However, more recent work of hypolithic communities have found support for the existence of dispersal limitation between deserts (Archer et al., 2019; Bahl et al., 2011; Caruso et al., 2011). Much of the work investigating lithic habitats over spatial scales has been done using the quartz hypolithic system [19,21,23,25] and only a few global surveys of the endolithic system exist (Friedmann, 1980; Walker and Pace, 2007). While hypolithic communities colonize the underside of quartz rocks at the interface with the underlying soil, endolithic communities colonize a diversity of rock substrates, have been found to be distinct from hypolithic communities within the same local environment (Van Goethem et al., 2016), and can even colonize more extreme environments (Warren-Rhodes et al., 2007a), raising the question of whether the same processes govern the assembly of both lithic habitats.

The central question this study seeks to answer is what factors determine the assembly of global desert endolithic communities. A variety of environmental effects have been proposed to influence endolithic community assembly, including macroclimate (Friedmann, 1980), rock geochemistry (Walker and Pace, 2007), and substrate architecture (Meslier et al., 2018), but study limitations and differing methodologies have made it difficult to place the relative importance of these factors. It also remains untested as to whether stochastic processes play a role in controlling endolithic community assembly. To investigate this question, we conducted a comprehensive survey of endolithic communities and relevant abiotic factors at global, continental, and regional scales. In order to focus on macroscopic influences, we only sampled sandstones to limit effects from the rock microenvironment, which have been shown to strongly affect community composition even within the same local environment (Crits-Christoph et al., 2016; Meslier et al., 2018).

## Materials and Methods

### Sample collection and DNA extraction

Colonized endolithic sandstones were collected from five desert regions (Negev Desert, Namib Desert, Colorado Plateau, Canadian Arctic, McMurdo Dry Valleys) between 2012 and 2018 (Supplementary Table S1). In the Namib Desert, Negev Desert, and Canadian Arctic, samples were collected from multiple regional sites (Fig. 1). Rock samples were broken off with a field hammer and stored in sterile WhirlPak bags (Nasco) in the dark at room temperature, except for samples from the Canadian Arctic and McMurdo Dry Valleys, which were stored in the dark at −20°C.

**Figure 1:**
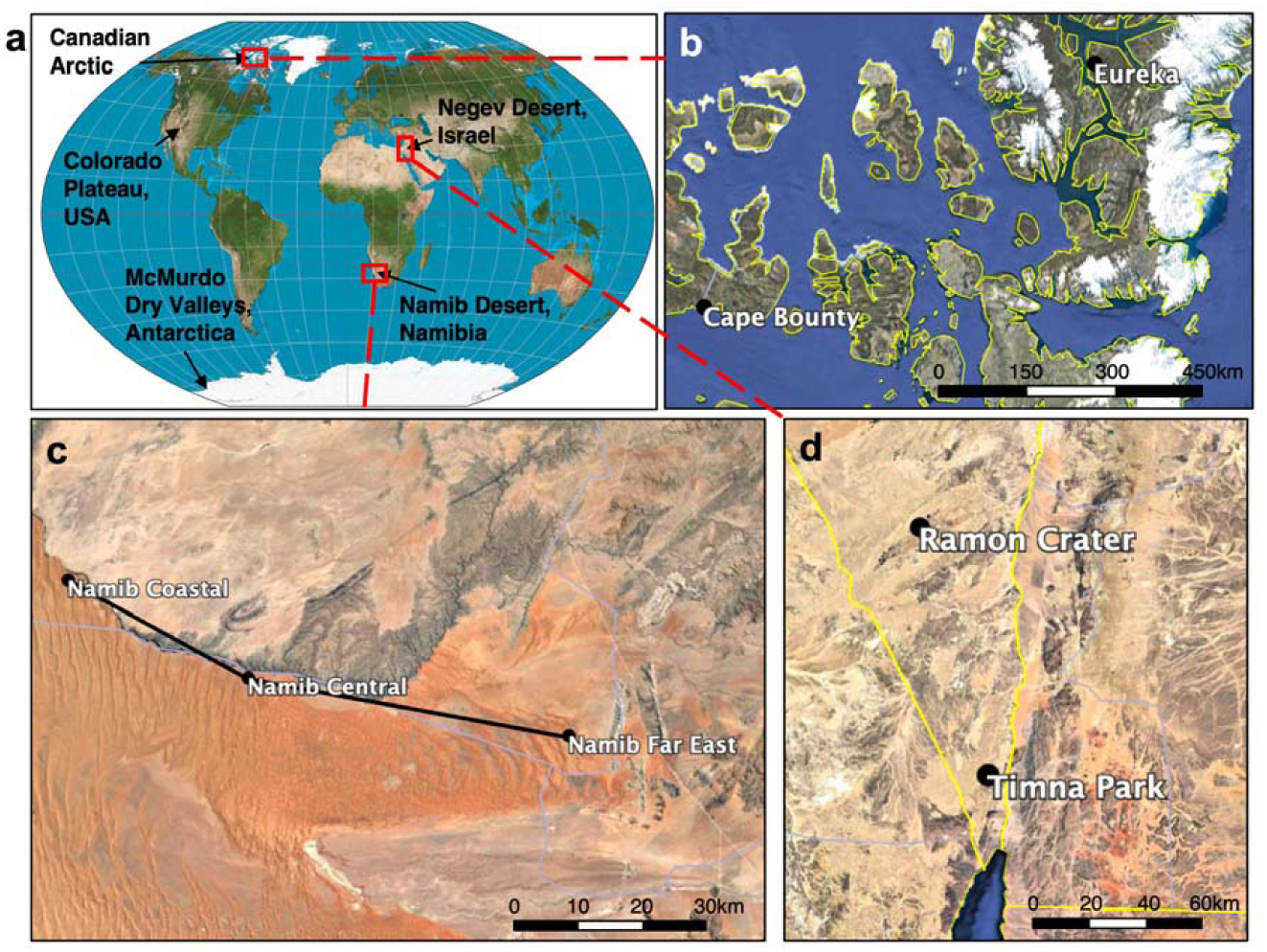
Map of global and regional sandstone sampling sites. (a) World map showing sampling locations. In the Canadian Arctic (b), Namib Desert (c), and Negev Desert (d) samples were taken from multiple sites whose names and locations are shown. More detailed information can be found in (Supplementary Table S1).

DNA was extracted from ten samples for each of the nine desert sites, for a total of 90 samples overall. Extraction was done with 0.25 ± 0.05g of rock scraped from colonized areas with a sterile metal tool using the PowerSoil DNA Isolation Kit (Qiagen). To validate the robustness of the sequencing pipeline, rock samples from each desert site were harvested and processed for sequencing in replicate. Post-extraction, DNA was quantitated with the Qubit Fluorometer dsDNA HS Assay (Thermo Fisher Scientific).

### Climate data

For each site, temperature, relative humidity, precipitation, and solar irradiance were obtained from meteorological sources for a two-year period encompassing the dates during which samples were collected (Supplementary Table S2). Daily solar radiation was found to be very consistent within desert sampled, and so a single average value was reported for each desert. In the Negev Desert, rainfall data were taken from 2017-2018, due to abnormal rainfall events in 2016 after sample collection that increased the two-year average. For the University Valley site, solar radiation data were obtained from a study of the nearby Beacon Valley.

### Scanning Electron Microscopy

Small colonized rock fragments were broken off from sandstones and fixed in 3.0% formaldehyde, 1.5% glutaraldehyde, 5mM Ca^2+^, 2.5% sucrose, in 0.1M sodium cacodylate, pH 7.4 at room temperature for 30 minutes. Afterwards, samples were washed thrice in 0.1M sodium cacodylate, 10 minutes each. Samples were stained with 1% OsO_4_ in 0.1M sodium cacodylate in the dark for 45 minutes, followed by three quick washes in ddH_2_O. Stained samples were dehydrated in a cold ethanol gradient: 50%, 75%, 90%, 95%, then three 15 minute changes in 100% ethanol at room temperature, followed by a final dehydration in hexamethyldisilazane for 2 minutes. Finally, samples were coated with a 1.5nm Pt layer before imaging in a FEI Quanta 200 Environmental SEM. Imaging was done at 15kV, spot size 1.5, 6mm working distance for high-magnification imaging and 20kV, spot size 2.5, 10mm working distance for low-magnification imaging.

### Measurement of physical and chemical sandstone properties

For each site, a combined 15g of crushed, non-colonized sandstone from three separate rocks was analyzed for mineral composition through borate fusion X-ray fluorescence, performed at SGS Canada Inc. (Lakefield, Canada). Sandstone samples were tested for soluble NO_3_^-^, PO_4_^3-^ Cl^-^, and SO_4_^2-^ via ion chromatography, EPA method 300.0 and soluble Ca^2+^, Mg^2+^, K^+^, Na^+^, Fe^2+^ and Fe^3+^ via inductively coupled plasma atomic emission spectroscopy, EPA method 200.7. Soluble ion analyses were performed at Inter-Mountain Labs (Sheridan, WY).

Grain size distribution of sandstone samples was obtained through sieve analysis (Krumbein and Pettijohn, 1938). Sandstones were first disaggregated by gently crushing 25g of rock with a pestle and mortar, taking care not to pulverize individual sandstone grains. Rock samples were shaken through a series of sieves with opening sizes 710, 500, 355, 250, 180, 125, 90, 60μm for 10 minutes, after which mass in each sieve was measured. Cumulative grain size distributions were fitted to a sigmoid function, from which a 50^th^ percentile diameter (D50) value was extracted for each rock. Water retention of sandstone rocks was measured through a fluid resaturation method. Rock samples in triplicate were first fully dried for 24h in an oven, then weighed. Samples were then immersed in distilled water for 3 days to allow the rock to fully saturate, after which wet mass and bulk volume (V_B_) were measured. Pore volume (V_P_) was obtained by subtracting dry mass from wet mass and multiplying by the density of water, and a percent water retention was calculated using the formula 100(*V*_*P*_ – *V*_*B*_) (U.S. Department of the Interior, 1963).

### 16S rRNA library generation and sequencing

Libraries for 16S rRNA gene and ITS amplicons were created using a two-step PCR protocol. The 16S rRNA gene was first amplified using primer pair 515F– GTGYCAGCMGCCGCGGTAA and 806R–GGACTACNVGGGTWTCTAAT targeting the V3-V4 hypervariable region (Caporaso et al., 2012). For ITS amplicon libraries, primer pair fits7F– GTGARTCATCGAATCTTTG and ITS4R – TCCTCCGCTTATTGATATGC was used (Ihrmark et al., 2012). The primer pairs ITS1F/ITS4R (Gardes and Bruns, 1993) and ITS86F/ITS4R (Turenne et al., 1999) were also tested on all samples to ensure eukaryotic diversity was thoroughly detected, but these did not lead to any additional amplification.

PCR reactions were constructed using: 4-40ng of template DNA, 1X Phusion PCR HF Buffer, 0.5mM MgCl_2_ 3% DMSO, 800µM dNTP mix, 0.25µM of each primer, 0.02U/µL of Phusion High-Fidelity DNA Polymerase (New England Biolabs), and nuclease-free water for a total volume of 25µL/reaction, and run with the following cycling conditions: 98°C for 30s, followed by 20 cycles of 98°C for 10s, 55°C for 15s, 72°C for 15s, final extension at 72°C for 5min. First step PCR reactions were done in duplicate for each sample, then pooled. After each PCR step, products were cleaned with a 0.6X Sera-Mag Speedbeads mixture (Roland and Reich, 2012), then quantitated with the Qubit Fluorometer dsDNA HS Assay (Thermo Fisher Scientific) before further downstream processing.

Barcode and adapter sequences were attached in a second step PCR reaction as follows: 4ng of cleaned first step PCR product, 1X Phusion PCR HF Buffer, 0.5mM MgCl_2_ 3% DMSO, 800µM dNTP mix, 0.36µM universal forward primer, 0.36µM uniquely barcoded reverse primer, 0.02U/µL of Phusion High-Fidelity DNA Polymerase (New England Biolabs), and nuclease-free water were combined for a total volume of 25µL/reaction. Cycling conditions for second step reactions were 98°C for 30s, followed by 10 cycles of 98°C for 10s, 55°C for 15s, 72°C for 15s, final extension at 72°C for 5min. Equimolar amounts of each library were pooled before sequencing, and final multiplexed libraries were quality-checked using the 2100 Bioanalyzer DNA High-Sensitivity Kit (Agilent) before sequencing. 2nM libraries were sequenced on the Illumina 2500 MiSeq platform at read length 2 x 250bp in the Genetic Resources Core Facility at Johns Hopkins University. Sequence data were demultiplexed and nonbiological sequences were removed prior to analysis.

### Analysis and visualization of sequence data

Eighty-nine 16S rRNA gene libraries and 29 ITS2 libraries from sandstone microbial communities were successfully sequenced, along with control replicates used to validate the robustness of the sequencing pipeline (Supplementary Table S1). Paired-end sequencing at 250bp returned 7,700,178 paired 16S rRNA gene reads and 2,070,702 paired ITS reads. Both bacterial and eukaryotic sequencing depths sampled community diversity to asymptote (Supplementary Fig. S1).

The bioinformatics package QIIME2, ver. 2018.11 was used to de-noise Illumina reads (Bolyen et al., 2018). Forward and reverse reads were manually trimmed at the 3’ end once the lower quartile of sequences dropped below a Phred score of 20. Quality filtering, dechimerization, and inference of exact sequence variants (ESVs) was done with the ‘dada2’ plugin in QIIME2 (Callahan et al., 2016). Sequencing runs were independently processed in the DADA2 pipeline and then consolidated. Due to high stochastic sequence variation at the ESV level, 16S rRNA gene analysis was performed after clustering ESVs into operational taxonomic units (OTUs). Using the ‘vsearch’ plugin in QIIME2 (Rognes et al., 2016) and the SILVA 16S rRNA reference database, release 128 (Quast et al., 2013), bacterial ESVs were clustered into open-reference OTUs at 97% similarity, returning 3227 OTUs and a 2.8-fold decrease in bins. ITS sequence analysis was done entirely at the ESV level due to the difficulties in accurately clustering ITS sequence data by identity (Popovic et al., 2018).

Taxonomy was assigned to eukaryotic ESVs and bacterial OTUs using a naïve-Bayes classification approach from the ‘q2-feature-classifier’ package in QIIME2 (Bokulich et al., 2018). The classifier was trained on the SILVA 16S rRNA reference database, release 128 at 97% identity (Quast et al., 2013) for bacterial sequences and the UNITE database ver. 7.2 at 97-99% dynamic identity for eukaryotic sequences (Nilsson et al., 2019). Mantel, adonis, and ANOSIM statistical tests, as well as Pearson correlation coefficient were calculated in QIIME2. Taxonomy, heatmap and PCoA visualizations were constructed in R using the ggplot2 and phyloseq packages. To generate bipartite networks, OTUs were first normalized using relative abundance, then node associations were calculated with the make_otu_network.py function in QIIME, release 1.9.1 (Caporaso et al., 2010). Network visualization was performed in Cytoscape, ver. 3.7.1 (Shannon et al., 2003).

## Results

Colonized endolithic sandstones were sampled in five deserts and nine sites (Fig. 1). Among deserts, a wide range of environmental conditions were observed (Fig. 2A-2E; Supplementary Table S2). We grouped these deserts into three distinct climate regimes based on mean temperature, solar irradiance, and moisture variables (Supplementary Table S2). Hot deserts (Namib and Negev Deserts) had high solar irradiance and air temperatures reaching up to 40°C in the summer months, as well as lower relative humidity. Polar deserts (Antarctic Dry Valleys and Canadian Arctic) were characterized by long winters with complete darkness and very low temperatures, followed by short summers with above-zero temperatures and near-constant sunlight. Temperate deserts (Colorado Plateau) was primarily characterized by high precipitation and relative humidity, along with moderate temperatures.

**Figure 2:**
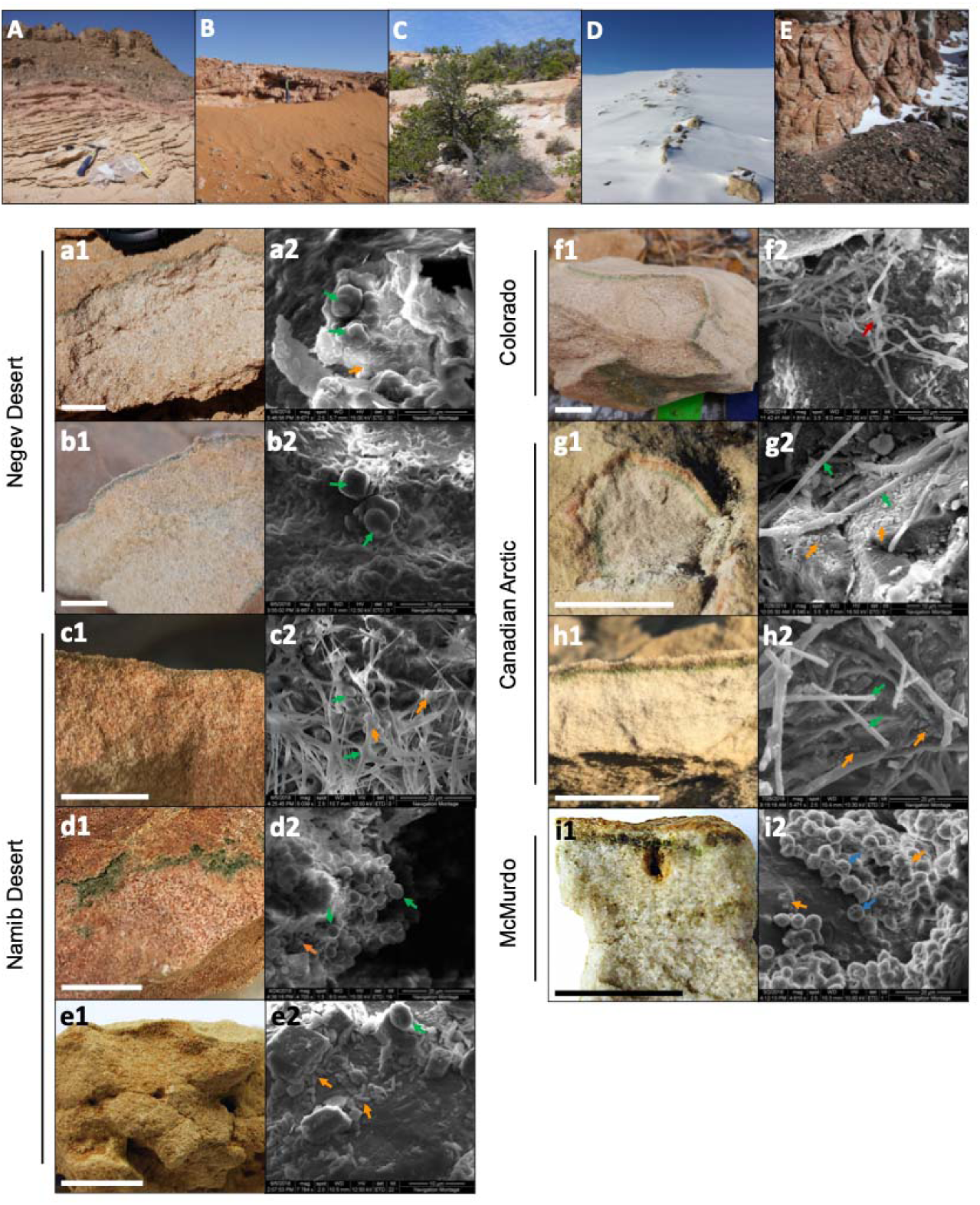
Location, colonization zone, and SEM micrographs of microbial communities from each sampling site. (**A-E**) General views of the 5 deserts sampled: (A) Negev Desert, (B) Namib Desert, (C) Colorado Plateau, (D) Canadian Arctic, and (E) McMurdo Dry Valleys. (**a1-i1**) Sandstone colonization zone at each site, in the order: (a) Timna Park, (b) Ramon Crater, (c) Namib Coastal, (d) Namib Central, (e) Namib Far East, (f) Escalante, (g) Eureka, (h) Cape Bounty, (i) University Valley. Scale bar = 2 cm. (**a2-i2**) SEM micrograph images of the endolithic microbial community. Green arrows indicate Cyanobacteria, orange arrows indicate heterotrophic bacteria, red arrows indicate fungal hyphae, and blue arrows indicate green algae.

### Characterization of endolithic colonization zones

In all sandstones, a green colonization zone was observed a few millimeters below the surface of the rock (Fig. 2a-i). The colonization was mostly cryptoendolithic with the exception of sandstone from central Namib, where colonization was found in small crevices parallel to the rock surface (chasmoendolithic; Fig. 2d). In Arctic and Antarctic sandstones, an additional orange (Fig. 2g) or brown (Fig. 2h-i) pigmented layer was located above the green colonization layer.

SEM imaging revealed microbial cells in dense aggregates within the colonized zone, with *Cyanobacteria* (Fig. 2a-h) and green algae (Fig. 2i) being the most prominent cells within the microbial landscape. Cyanobacterial morphologies varied between deserts and sites, ranging from mostly coccoidal in the Negev Desert (Fig. 2a-b) to filamentous in the Canadian Arctic (Fig. 2f-g). Coccoidal cyanobacteria had a diameter of 3-4 μm, while filamentous *Cyanobacteria* were around 1 μm in diameter and up to 50 μm long. Green algae, 5-6 μm in diameter, were observed only in the Antarctic University Valley community. Coccoidal and bacilliform heterotrophic bacteria were found associated with phototrophs in all communities; their sizes ranged between 0.5 and 1 μm in the longest dimension. Large fungal hyphae measuring up to 20 μm wide and over 200 μm long were observed in the Colorado Plateau community (Fig. 2h). Microbial cells were often found surrounded by extracellular polymeric substances (EPS), with the organization of EPS varying greatly between deserts and sites. For example, in Negev Desert samples, *Cyanobacteria* and heterotrophic bacteria were organized in clusters and encased entirely in EPS (Fig. 2a-b).

### Rock architecture and chemical properties

X-ray fluorescence (XRF) analysis showed that SiO_2_ was the most abundant compound in all sandstones, followed by Fe_2_O_3_ and Al_2_O_3_. The compounds CaO, MgO, K_2_O, Na_2_O, TiO_2_, and MnO were present in all sandstones at abundances less than 2%, while P_2_O_5_, Cr_2_O_3_, and V_2_O_5_ were sparsely present in sandstones at abundances less than 0.1%. (Supplementary Table S3). Namib Central sandstones had a unique XRF profile with high percentages of CaO (15.8%) and loss on ignition mass (13.5%), suggesting an enrichment in calcium carbonate, which breaks down into CaO and O_2_ during analysis (Supplementary Table S3). Water-soluble ions were measured by ion chromatography and inductively coupled plasma atomic emission spectroscopy and remained at concentrations below 30 mg/kg. Soluble Fe^2+^, Fe^3+^, and PO_4_^3-^ were below detection limits (Supplementary Table S4). Sandstone grain size was variable between sites, with the D50 of sandstone grains ranging from 116 to 530μm (Supplementary Table S5, Supplementary Fig. S2). Water retention capacity was measured with a fluid resaturation method and ranged from 10.9% to 28.0% v/v (Supplementary Table S5).

### Endolithic community composition

High-throughput sequencing of the 16S rRNA gene and ITS regions was used to characterize sandstone endolithic communities at the molecular level. Bacteria and Eukarya were the most abundant members of the endolithic community, while Archaea were only present at very low abundances (<0.1%) in every desert except the McMurdo Dry Valleys (Fig. 3a). The most abundant bacterial phyla were *Cyanobacteria, Actinobacteria*, and *Proteobacteria*, which together represented 48-75% of the communities (Fig. 3b). The eukaryotic community was comprised of fungi belonging to the phyla *Ascomycota* and *Basidiomycota* along with the green algae *Trebouxia* (Fig. 3c). Archaea primarily belonged to the phylum *Thaumarchaeota*. Relative abundance of eukaryotes within the overall microbial community was roughly estimated from the number of mitochondrial (mt) and chloroplast (cp) DNA reads in the 16S rRNA gene sequencing data (Fig. 3a). mtDNA and cpDNA reads were only detected in the polar deserts (University Valley, Eureka, Cape Bounty) and to a lower extent, in the temperate Escalante community. These results were supported by ITS sequencing, for which amplicons could only be obtained for the Colorado Plateau, Canadian Arctic, and Antarctic Dry Valley communities (Fig. 3c). The taxonomic assignment of fungal ITS sequences performed particularly poorly, with 44% of ITS sequence variants only assignable to the phylum level or higher.

**Figure 3:**
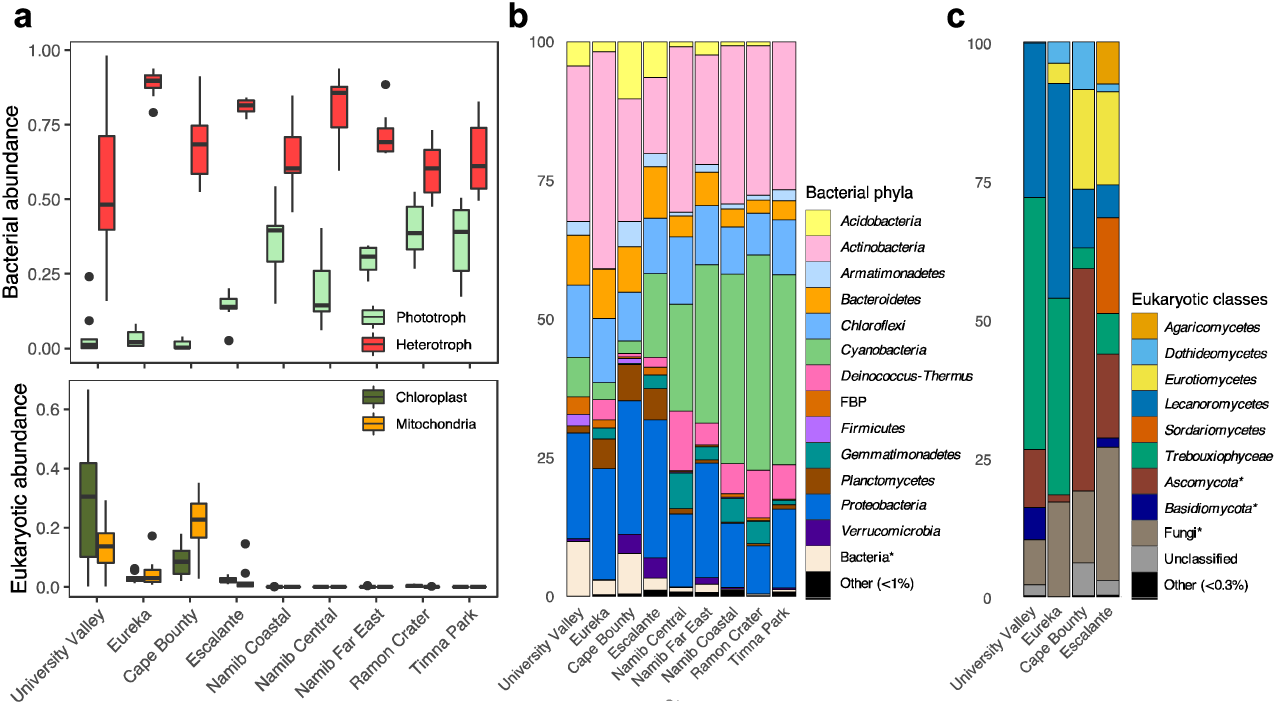
Sandstone endolithic community composition. (**a**) Top: Relative abundances of bacterial prokaryotes and eukaryotes, obtained from 16S rRNA gene sequences Bottom: Relative abundances of chloroplast and mitochondrial 16S rRNA gene sequences (**b**) Prokaryotic community composition, based on 16S rRNA gene sequences. (**c**) Eukaryotic community composition based on ITS sequences. Asterisks indicate a lack of taxonomic information up to the designated level.

Phototrophs, consisting of *Cyanobacteria* and the green alga *Trebouxia*, comprised 21 ± 17% and 18 ± 23% of the bacterial and eukaryotic community, respectively (Fig. 3b, 3c). There was a significant negative relationship between *Cyanobacteria* and algae relative abundances (Pearson; ρ = −0.539, p < 0.001). This was especially true for the University Valley and Eureka sites, where *Trebouxia* was highly abundant and few *Cyanobacteria* were detected (Fig. 3).

### Global effects of endolithic community assembly

Initial patterns of bacterial community assembly were analyzed through principal coordinates analysis (PCoA) of weighted UniFrac distance (Fig. 4a). Communities significantly clustered according to their respective climate regime (ANOSIM; R = 0.87, p < 0.001); this sample grouping alone explained 46% percent of the variation between communities (adonis; R = 0.46, p < 0.001). Among individual climate variables, average air temperature was the most correlated with community distance, followed by yearly solar irradiance and distance between sampling sites (Fig. 4b, Table 1). Only a few substrate variables were significantly correlated with community distance: soluble K^+^, percent Fe_2_O_3_, and percent Al_2_O_3_ (Table 1).

**Table 1:**
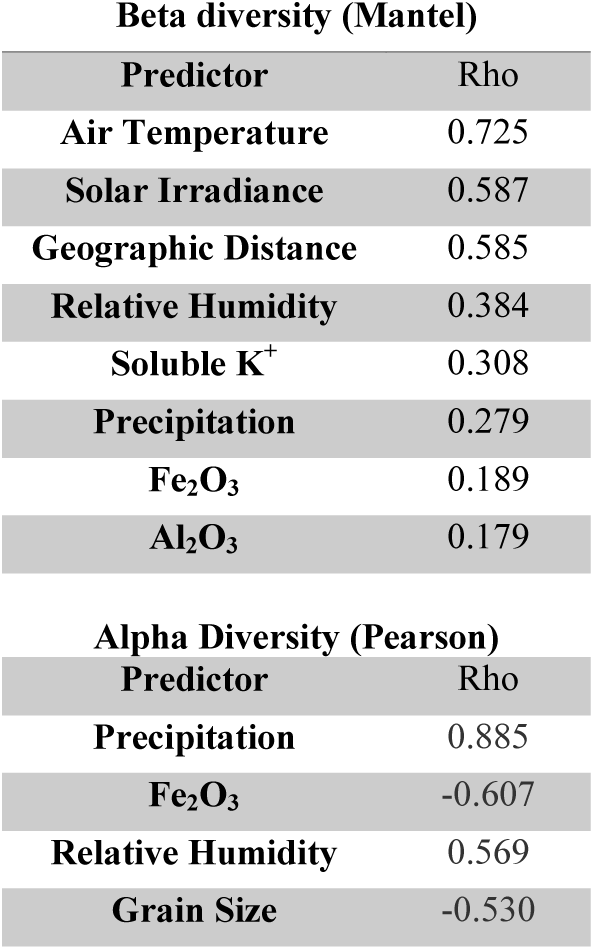
Significant correlations (p<0.01) of abiotic variables with community alpha and beta diversity, using Pearson and Mantel tests.

| Predictor | Rho |
| --- | --- |
| Air Temperature | 0.725 |
| Solar Irradiance | 0.587 |
| Geographic Distance | 0.585 |
| Relative Humidity | 0.384 |
| Soluble $K^+$ | 0.308 |
| Precipitation | 0.279 |
| $Fe_2O_3$ | 0.189 |
| $Al_2O_3$ | 0.179 |

**Alpha Diversity (Pearson)**
| Predictor | Rho |
| --- | --- |
| Precipitation | 0.885 |
| $Fe_2O_3$ | -0.607 |
| Relative Humidity | 0.569 |
| Grain Size | -0.530 |

**Figure 4:**
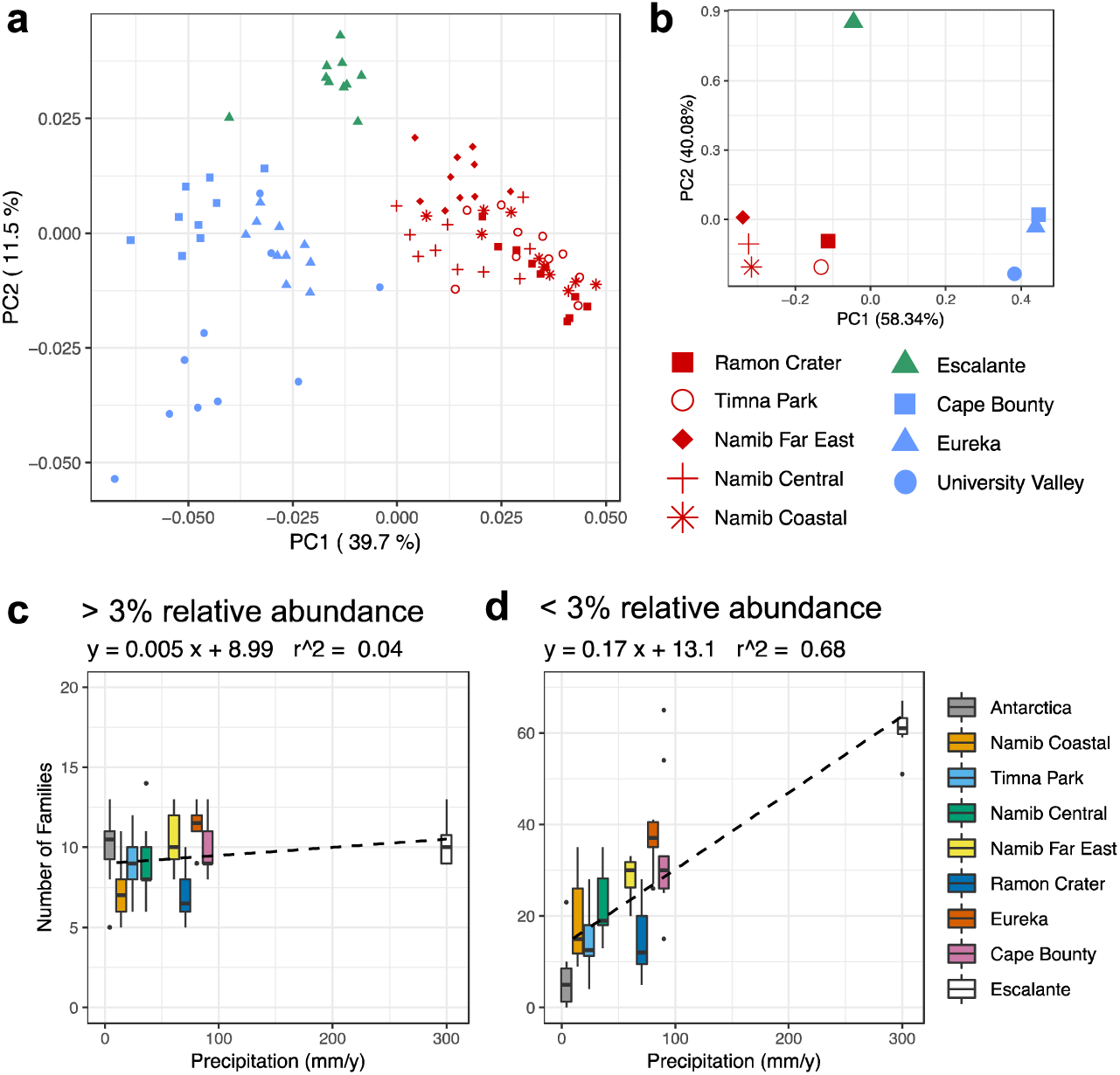
Environmental correlations with bacterial community composition and diversity. (**a**) Principal coordinates analysis (PCoA) plot of weighted UniFrac distance of the prokaryotic community, based on 16S rRNA gene sequences rarefied to 1500 reads. Sites are colored according to climate regime, with red = hot/dry, blue = polar, and green = temperate. (**b**) PCA of Euclidean distance of climate variables: average temperature, precipitation, relative humidity, daily solar radiation. (**c-d**) Boxplots of number of families plotted against yearly precipitation for (**c**) highly abundant OTUs and (**b**) lowly abundant OTUs.

Precipitation was the variable most correlated with bacterial and eukaryotic alpha diversity; grain size, relative humidity and percent Fe_2_O_3_ were also significantly correlated (Table 2). The community response to precipitation was remarkably different for highly and lowly abundant taxa, with the strongest effect evident at the family level (Fig. 4d, 4e). The endolithic community harbored 7-13 highly abundant families, (> 3% relative abundance) independently of moisture conditions (R^2^ = 0.04), but the number of lowly abundant families (< 3% relative abundance) increased in a positive linear relationship (R^2^ = 0.68) from around 5 families at the driest site (University Valley) to over 60 at the wettest (Escalante).

### Fine-scale partitioning of the sandstone microbial community

In order to identify which taxa drive community differences between climate regimes, bipartite networks were generated for the three most abundant bacterial phyla: *Cyanobacteria, Actinobacteria*, and *Proteobacteria* (Fig. 5). Network analysis revealed that *Cyanobacteria* operational taxonomic units (OTUs) were particularly specific to a single climate regime, with only 18% of cyanobacterial OTUs shared between two or more climate regimes (Fig. 5a, Supplementary Fig. S3). Hot deserts (Fig. 5a, red boxes) had a cyanobacterial profile that was dominated by *Chroococcidiopsis*, with a single *Chroococcidiopsis* OTU (accession number FJ790616) highly abundant in all hot communities. Polar deserts (Fig. 5a, blue boxes) had the lowest cyanobacterial diversity among the climate regimes, especially at University Valley (UV) and Eureka (EU). *Chroococcidiopsis, Cyanothece*, and *Phormidium* were the most common *Cyanobacteria* found in polar deserts. The highest cyanobacterial diversity was located in the temperate Escalante (ES) site (Fig 5a, green boxes), which uniquely harbored 47% of cyanobacterial OTUs and had higher taxonomic diversity than other deserts. The wetter Namib Far East (FE) site also had higher OTU and taxonomic diversity, albeit to a lesser extent.

**Figure 5:**
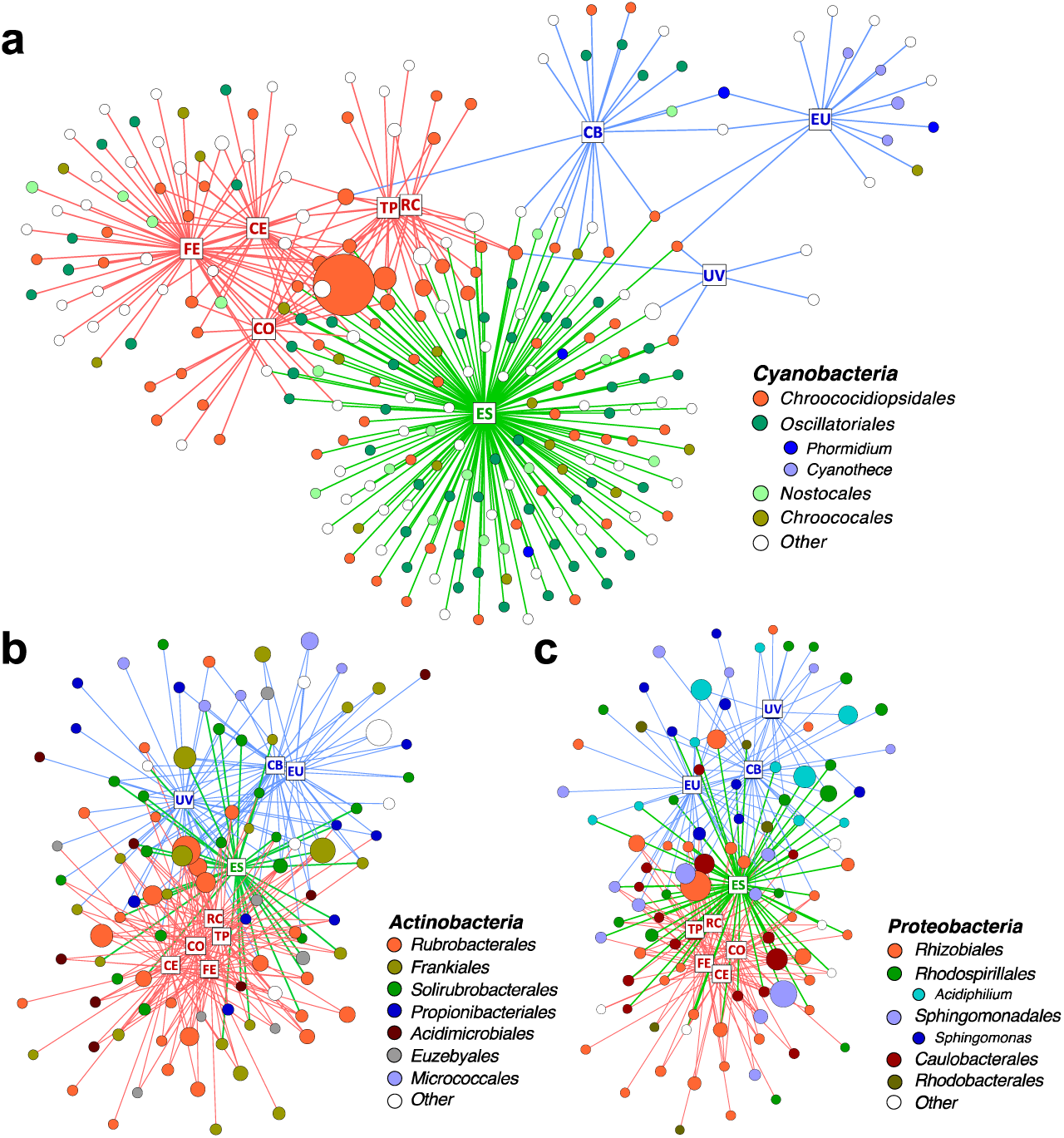
Distribution of OTUs across desert sites. Bipartite network of (**a**) all *Cyanobacteria* and 100 most abundant (**b**) *Actinobacteria* and (**c**) *Proteobacteria* OTUs. Squares represent sites and are labeled with site name: RC = Ramon Crater, TP = Timna Park, CO = Namib Coastal, CE = Namib Central, FE = Namib Far East, ES = Escalante, CB = Cape Bounty, EU = Eureka, UV = University Valley. Edges are colored in according to the climate regime of their respective site: red = hot, blue = polar, and green = temperate. Circles represent OTUs and are colored by taxonomic order and genus. Singleton taxa and unassigned OTUs are colored in white. The size of each circle is proportional to mean relative abundance.

Heterotroph OTUs were distributed in a more cosmopolitan manner, with 54% of *Actinobacteria* OTUs (Fig. 5b, Supplementary Fig. S3) and 65% of Proteobacteria OTUs (Fig. 5c, Supplementary Fig. S3) shared between two or more climate regimes. OTU richness among *Proteobacteria* and *Actinobacteria* did not vary between climate regimes, but differences in taxonomic distribution were noticed between regimes. For instance, 34% of *Actinobacteria* OTUs in hot deserts belonged to the single genus *Rubrobacter* and the proteobacterial genera *Acidiphilum* and *Sphingomonas* were mostly specific to polar deserts.

### Regional impacts on endolithic community assembly

Sandstones were collected at multiple sites within the Canadian Arctic, Namib Desert, and Negev Desert to further investigate within-desert impacts on community assembly (Fig. 1). Significant community differences were observed between sites in the Canadian Arctic and Namib Desert, but not the Negev Desert. In the Canadian Arctic, the eukaryotic community at Eureka was low-diversity, containing 29 ± 10 exact sequence variants (ESVs) and dominated by *Trebouxia* (36%), while the eukaryotic community at Cape Bounty was high-diversity, containing 74 ± 14 ESVs and *Trebouxia* was lowly abundant (4%) (Fig. 1, Supplementary Table S6). Within the Namib Desert, hierarchical clustering analysis at the OTU level showed that bacterial profiles generally clustered according to site, but this grouping was much more rigorous for *cyanobacteria* than for heterotrophic bacteria (Fig. 6, Fig S4). Notably, the Namib Far East site harbored a much higher *Cyanobacteria*l diversity than the other two Namib sites, but not a considerably higher heterotrophic diversity.

**Figure 6:**
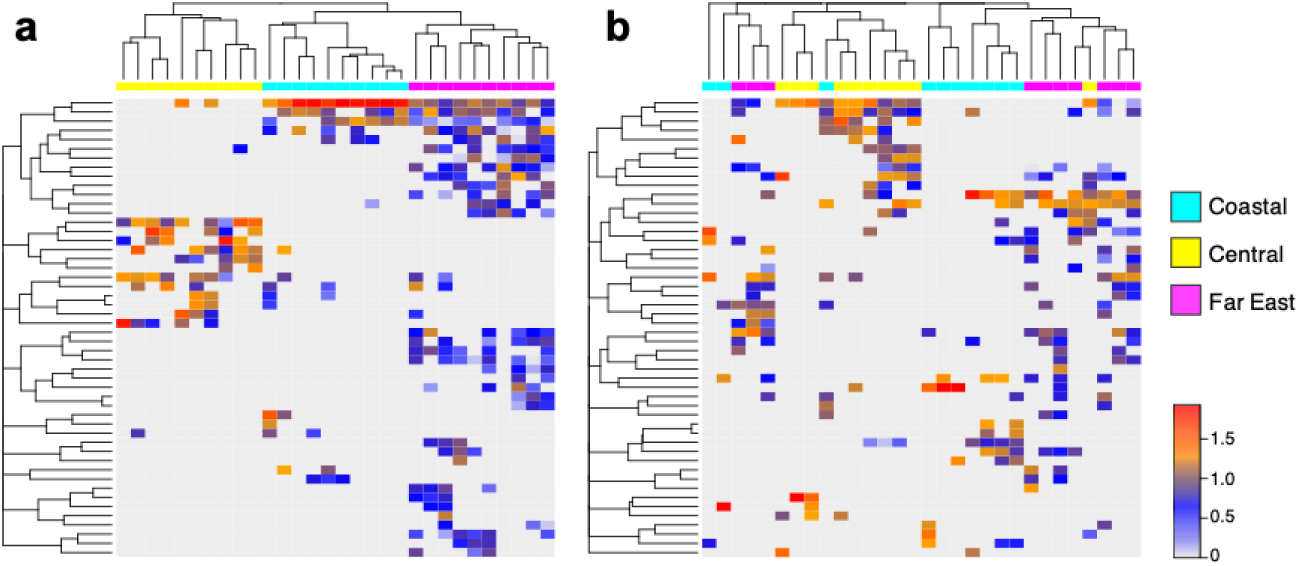
Regional differences in Namib Desert communities. UPGMA-clustered heatmap of 50 most abundant (**a**) *Cyanobacteria* OTUs and (**b**) *Proteobacteria* OTUs from three sites in the Namib Desert. Rows correspond to OTUs and columns correspond to individual samples. Color scale represents log-normalized relative abundances.

## Discussion

We sampled endolithic sandstones across three major biogeographic regions (North America, Africa/Asia, and Antarctica) and three diverse climate regimes in order to investigate long-range patterns of diversity in endolithic communities. Previous global studies of the endolithic system only reported samples from 2 regions, using low-resolution molecular analysis (Friedmann, 1980; Walker and Pace, 2007). Our sandstone samples had mostly similar geochemical properties, thus confirming that the sandstone endolithic system is a consistent model for studying macroscopic patterns in microbial ecology.

The bacterial composition of the sandstone community was consistent with previous findings from sandstones (Walker and Pace, 2007), as well as other endolithic substrates (Meslier et al., 2018). The ubiquitous phyla in nearly all endolithic substrates were *Cyanobacteria, Proteobacteria*, and *Actinobacteria*, while the phyla *Acidobacteria, Deinococcus-Thermus, Bacteroidetes, Chloroflexi*, and *Planctomycetes* were commonly found but often less abundant. The eukaryotic component of the endolithic community has been more poorly characterized, and the sparse taxonomic annotation of ITS sequences in this study suggests that much of the eukaryotic diversity within endoliths remains to be discovered. Despite this, our findings concur with previous studies which report that the eukaryotic community is comprised of lichenized fungi from the phyla *Ascomycota* and *Basidiomycota* along with green algae from the phylum *Chlorophyta* (Archer et al., 2017; Pointing et al., 2009). While we were only able to find eukaryotes within polar and temperate desert samples, this does not necessarily indicate that no eukaryotes are present in hot desert samples and may instead reflect primer biases associated with ITS-based eukaryotic profiling (Popovic et al., 2018). Archaea have been poorly reported in endolithic communities to the point where their presence is questioned (Meslier et al., 2018; Pointing et al., 2009); however, we found archaeal signatures in every site except University Valley, indicating that Archaea do contribute to the sandstone endolithic community, albeit at very low relative abundances.

We did not find support for geographic distance alone being a primary driver of community assembly, and it is likely that the biogeographic signal in our data is mostly an artifact of environmental differences between continents. The most convincing evidence for this is the remarkable similarity in community composition between the Arctic and Antarctic communities and the bipolar distribution of many taxa, a finding which has been replicated in soil (Cox et al., 2016) and biofilm communities (Kleinteich et al., 2017). Highly similar communities were also found between the hot deserts; these results together suggest that environmental filtering, rather than dispersal limitation, is the primary organizing principle in endolithic communities. While previous studies have used similar findings as evidence for global dispersal of microorganisms throughout the atmosphere (Walker and Pace, 2007), a preponderance of recent evidence has been found in support of widespread dispersal limitation among microbial communities (Archer et al., 2019; Moeller et al., 2017; Power et al., 2018). In Antarctica, the high aerial environment has been shown to be particularly selective and limits dispersal of many microbial taxa to and from the continent (Archer et al., 2019). This raises the possibility that the bipolar distribution of certain OTUs may be a remnant of ancient biogeography (Bahl et al., 2011).

We pinpoint macroclimate as the primary driver of global endolithic community assembly. The greatest difference in community composition was observed between polar and hot desert conditions. The hot desert communities were almost entirely bacterial and dominated by *Cyanobacteria*. In contrast, communities from polar deserts had greatly reduced relative abundances of *Cyanobacteria* and saw an increased presence of fungi and green algae. The temperate Colorado community supported rich *Cyanobacteria*l and eukaryotic populations and was the most diverse site by far. This difference in community composition between hot and cold deserts has been well-studied by Friedmann et al. (Friedmann, 1980), who first posited that conditions in hot deserts were actually harsher for life than in polar deserts. In hot deserts, high daytime evapotranspiration causes the endolithic habitat to be desiccated for a majority of the time, with only a short period of time in the early morning suitable for photosynthesis (Friedmann, 1980). Conversely, in polar deserts humidity and constant solar radiation during summer months creates a relatively warm and wet environment in the rock matrix in spite of below-freezing outside conditions (Omelon et al., 2006). Thus, the patterns of global community composition are likely a result of eukaryotic life being mostly excluded from the harsher hot deserts, with the reappearance of eukaryotes in the more temperate Escalante site supporting this hypothesis. A similar pattern has been described regionally in halite nodules from the Atacama Desert, Chile, where algae were only found in the less hyper-arid areas of the desert (Robinson et al., 2015).

Moisture availability has generally been assumed to be the most important factor for desert communities under constant xeric stress (Lebre et al., 2017; Neilson et al., 2017; Pointing et al., 2007); however, we found that moisture variables such as precipitation and relative humidity are secondary drivers of global community assembly, behind hot versus polar conditions. The most noticeable community response to higher moisture was an increase in the number of low abundance taxa, including many non-desert adapted taxa. These lowly abundant taxa were often desert-specific, supporting the concept that stochastic assembly gains increased importance at higher moisture regimes (Chase, 2010; Lee et al., 2018). On the other hand, the composition and diversity of highly abundant and presumably well-adapted taxa were found to be remarkably similar across moisture regimes, particularly at the family taxonomic level. This is in contrast with desert soil communities, where major reorganization from drier to wetter conditions have been reported (Lee et al., 2018; Neilson et al., 2017; Scola et al., 2018). The taxonomic consistency across various moisture regimes suggests that a core community exists with a functional landscape that is well-adapted to the rocky endolithic habitat, regardless of moisture regime. We should note that this observation is not necessarily true when comparing rock substrates of different compositions and physico-chemical properties, as significant differences were observed in the composition and diversity of communities from different types of substrates found side-by-side in the Atacama Desert (Meslier et al., 2018).

Sandstone rocks have been found to support cryptoendolithic communities (Walker and Pace, 2007); however, we also found a unique chasmoendolithic community in the calcium carbonate-rich Namib Central sandstone, supporting a cyanobacterial profile not found in other Namib sandstones. Both the chasmoendolithic and cryptoendolithic Namib sandstone were still dominated by the genus *Chroococcidiopsis*, but they harbored substrate-specific OTUs, suggesting specialization of closely related species depending on substrate properties. Fine-scale partitioning of microbial diversity by substrate type has been described in calcite, ignimbrite and gypsum rocks and mainly attributed to “rock architecture”, meaning the space availability and water retention properties of the rock matrix (Crits-Christoph et al., 2016; Meslier et al., 2018). Other properties of the rock substrate, such as geochemical composition and soluble ion availability were generally not found to influence microbial assembly, with the exception of Fe_2_O_3_ and Al_2_O_3_ abundances. While these minerals might directly influence community physiology, they might also impact community assembly by affecting light transmission properties of the rock (McKay, 2012).

We found that at both global and regional scales, microbial community assembly partitioned by trophic level. Globally, phototrophic taxa were highly exclusive to their respective climate regime while heterotrophic taxa had a more cosmopolitan distribution. Regional inputs such as substrate properties and local moisture gradients also affected phototrophs more than heterotrophs. We argue these results demonstrate differential selective pressures between producers and consumers that drive endolithic community assembly.

Endolithic producers are likely more sensitive to climatic conditions such as water availability, solar irradiance, and temperature due to their photosynthetic requirement and status as pioneering organisms in endolithic communities (Crispim and Gaylarde, 2005; Tiano et al., 1995). In hyper-arid and hot deserts, *Chroococcidiopsis* is the only phototroph found in the community; this has been attributed to the superlative desiccation and radiation resistance properties of the taxon (Billi et al., 2000, 2011). It has been argued that the combination of hot and hyper-arid conditions constitutes the most extreme terrestrial environment on Earth (Friedmann, 1980; Wierzchos et al., 2018), suggesting that the absence of other types of phototrophs in the most extreme deserts is likely the result of stringent environmental filtering. Indeed, even a modest increase in precipitation within the Namib Desert resulted in a large increase in the diversity of *Cyanobacteria*. In contrast, polar communities were dominated by eukaryotic *Trebouxia* while *Cyanobacteria* were only sparsely present, suggesting that prokaryotic and eukaryotic phototrophs compete for access to the limited space within the rock suitable for photosynthesis. This is supported by the high number of secondary metabolites, especially antimicrobial compounds, that are found in endolithic *Cyanobacteria* (Crits-Christoph et al., 2016). Furthermore, *Trebouxia*, as well as the fungal classes *Lecanoromycetes* and *Eurotiomycetes* are well-recognized for their tendency to form lichen symbionts (Ahmadjian, 1988; Geiser et al., 2008; Miadlikowska et al., 2006) and lichen are particularly well adapted to the polar environments because they are more cold-resistant than the photobiont or mycobiont components alone (Barták et al., 2007). Physiological studies also show that endolithic lichens begin to photosynthesize at lower water potentials than *Chroococcidiopsis*, allowing them to better utilize the abundant relative humidity in polar deserts (Palmer and Friedmann, 1990). High-resolution sequencing of the semi-arid Escalante site revealed a scale of diversity previously undescribed in desert endolithic systems, with a community supporting robust prokaryotic and eukaryotic phototrophic populations. It has been suggested that local geochemistry, in particular pH, plays a more important role in driving community structure under milder conditions (Walker and Pace, 2007).

Heterotrophs exhibited higher stochasticity across deserts at both a global and regional scale. Examination of heterotrophic taxa across the Namib Desert moisture gradient found little difference in diversity, suggesting that consumer assembly is less sensitive to climate-based selection than producer assembly. Features of some heterotrophic taxa, such as desiccation resistance and spore formation may also promote their long-range dispersal in the atmosphere and contribute to the more cosmopolitan global distribution we observed (Archer et al., 2019). Despite this, network analysis showed that heterotrophic taxa still tended to cluster according to their respective climate regime, reinforcing the view that consumers still remain organized according to environmental influences. However, the patterns of consumer assembly we observed were more difficult to explain than those of producer assembly. One likely explanation is that biotic influences play a significant role in driving consumer profiles across deserts, including specific interactions with producers and predation from viruses (Fernández et al., 2018; Rodriguez-Brito et al., 2010; Valverde et al., 2015), but these dynamics are poorly studied in the endolithic system.

The partitioning of assembly influences by trophic level has been proposed before in studies of the hypolithic system but these studies do not concur on how phototroph and heterotroph assemblies are different (Caruso et al., 2011; Lacap-Bugler et al., 2017). We also note that these previous studies reported modest assembly according to environmental niche and the reported effects were small when compared to the strong climate-based clustering we observed within the endolithic system. Hypolithic communities are contiguous with the underlying soil and recruited in part from local soil populations (Makhalanyane et al., 2013), which may explain the increased stochasticity observed in hypolithic community assembly. In contrast, endolithic communities are completely enclosed within the rock environment and are further influenced by unique properties of the rock substrate, such as geochemistry, water retention, and light attenuation (Meslier et al., 2018). We suggest that complex assembly processes may be more discernible in the endolithic system because the assembly of these communities is influenced by more tractable environmental variables; thus the endolithic system may prove promising for further investigations of microbial biogeography.

## Conclusions

This study characterized for the first time global patterns of endolithic community diversity using high-throughput sequencing. We found that climate-based environmental selection was the primary organizer of endolithic community composition. This study joins several others in demonstrating that desert microbiomes are highly sensitive to climatic conditions, highlighting a need for studies of these fragile communities in the context of changing climate. The importance of rock substrate properties in the endolithic community is also evident, and we found a unique chasmoendolithic sandstone community in the Namib Desert, which demonstrates that within a single desert, substrate effects may be equally or more important than microclimate in organizing community composition. This study also reveals that phototrophs and heterotrophs within the endolithic community organize according to different assembly influences, with phototrophs being more sensitive to climate and substrate effects than heterotrophs. These findings emphasize that the functional niche is an essential consideration in studies of microbial assembly, and future research on lithic systems should consider using metaomic methods to investigate community function along with taxonomy.

## Data Availability

Raw sequencing data are available from the National Center for Biotechnology Information Sequence Read Archive (SRA) under BioProject ID PRJNA561762.

## Author Contributions

EQ contributed to the design of the study, conducted the experiments, analyzed the data, and wrote the manuscript. JDR, AO, and DC collected field samples. GMK provided support for fieldwork in the Namib Desert. CO collected field samples and took images from the Canadian Arctic. VM and JDR conceived and contributed to the design of the study. JDR supervised the project and edited the manuscript. All authors approved the final version for submission.

## Acknowledgments

We thank the entire staff of Timna Park for their kind hospitality during our field expedition. We also thank the staff of the Gobabeb Research and Training Centre (Namibia) for their support in the field, as well as the Cape Bounty Arctic Watershed Observatory and Environment Canada’s Eureka Weather Station for logistical support. We also acknowledge the technical support of Michael McCaffery for microscopy analysis. This work was supported by NSF grant DEB1556574.

