## Supplemental Materials for "Trophic Selective Pressures Organize the Composition of Endolithic Microbial Communities from Global Deserts"

**Supplementary Materials**

Evan Qu^1^, Chris Omelon^2^, Aharon Oren^3^, Victoria Meslier^1^, Don A. Cowan^4^, Gillian Maggs-Kölling^5^, and Jocelyne DiRuggiero^1§^

^1^*Johns Hopkins University, Department of Biology, Baltimore, USA*

^2^*McGill University, Department of Biology, Montreal, Quebec Canada*

*^3^The Hebrew University of Jerusalem, Institute of Life Sciences, Edmond J. Safra Campus, Jerusalem, Israel*

*^4^Centre for Microbial Ecology and Genomics, Department of Biochemistry, Genetics and Microbiology, University of Pretoria, Pretoria, South Africa*

*^5^Gobabeb-Namib Research Institute, Walvis Bay, Namibia*

**Table of Contents**

**Table S1:** Sampling sites and dates, number of rocks sequenced per site

**Table S2:** Weather data from sites.

**Table S3:** Chemical composition of sandstone samples

**Table S4:** Water-soluble ions from sandstone samples

**Table S5:** Sandstone physical properties

**Table S6:** Alpha diversity by site

**Figure S1:** Diversity rarefaction curves

**Figure S2:** Grain size distributions for sandstones

**Figure S3:** Venn diagrams showing distribution of bacterial OTUs

**Figure S4:** Heatmap of Namib Desert Actinobacteria

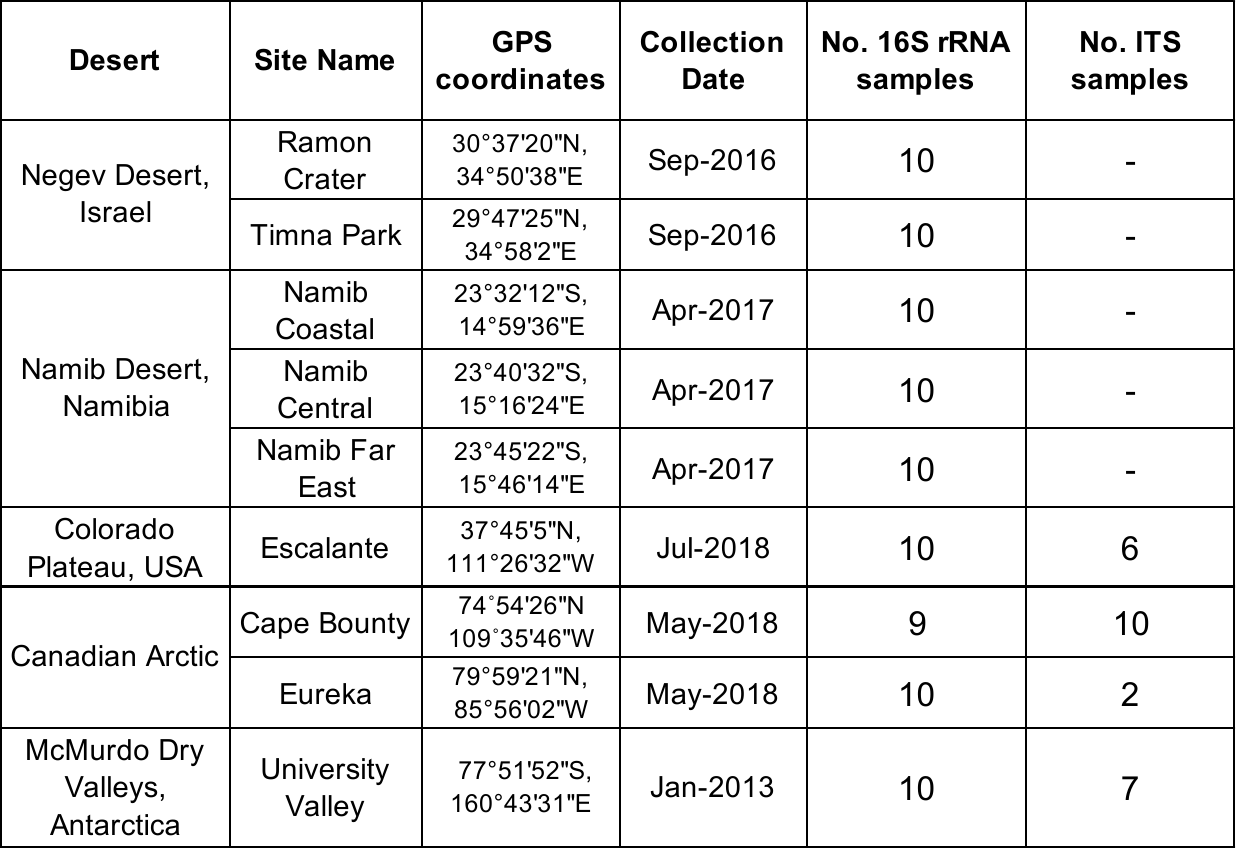

**Table S1: Sampling sites and dates, number of rocks sequenced per site.**

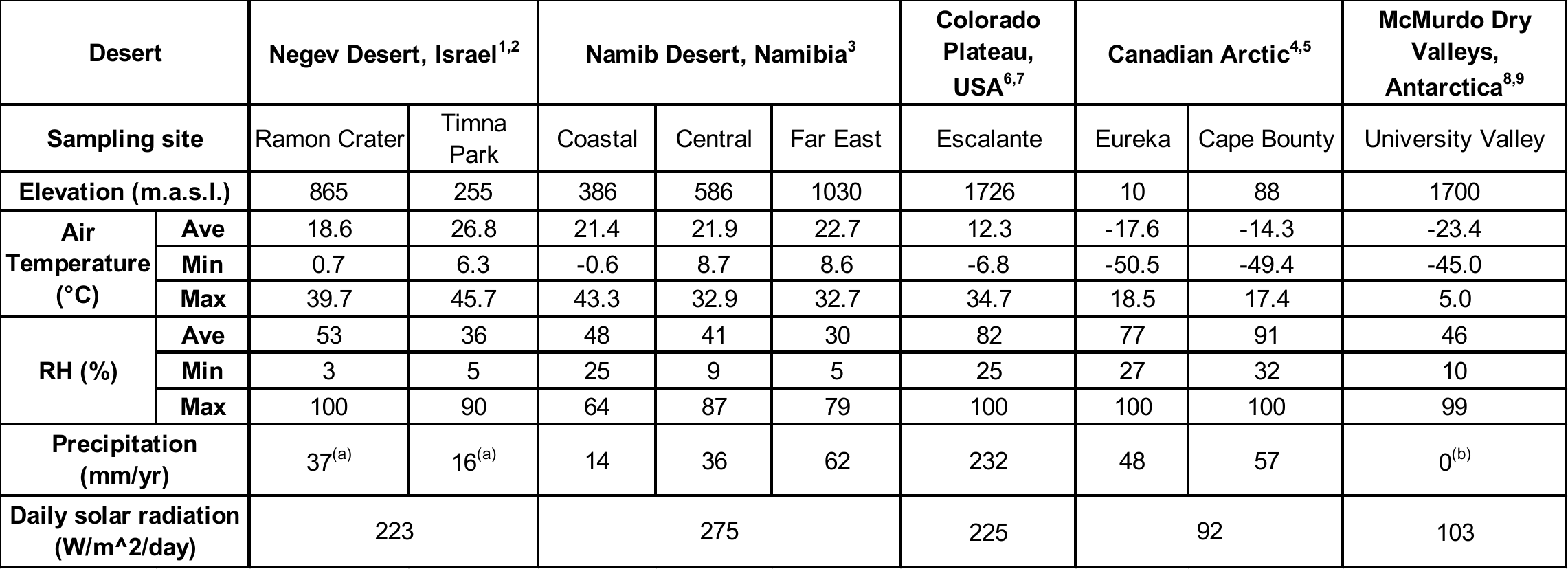

^1^Israel Meteorlogical Service - Mitzpe Ramon and Eilat stations; <http://www.ims.gov.il> **|** 2015-7 to 2017-7

^2^Evseev and Kudish, Analysis of solar irradiation measurements at Beer Sheva, Israel from 1985 through 2013, *Energy Conservation and Management*. 97: 307-314 (2015) **|** 2012-1 to 2013-8

^3^Sasscal WeatherNet - Gobabeb Met, Garnet Koppie, Ganab stations; <http://www.sasscalweathernet.org> **|** 2015-7 to 2017-7

^4^Environment and Natural Resources, Government of Canada **|** 2016-1 to 2017-12

^5^Cape Bounty Arctic Watershed Observatory; <https://capebountyresearch.com/> **|** 2016-1 to 2017-12

^6^NOAA National Centers for Environmental Information - Escalante and Bryce Canyon stations; <https://www.ncdc.noaa.gov> **|** 2017-1 to 2018-12

^7^NOAA National Solar Radiation Database - Bryce Canyon Airport station; <https://www.ncdc.noaa.gov> **|** 2010-1 to 2010-12

^8^Lacelle et al, Solar Radiation and Air and Ground Temperature Relations in the Cold and Hyper-Arid Quartermain Mountains, McMurdo Dry Valleys of Antarctica, *Permafrost and Periglac. Process*. 27: 163–176 (2016) **|** 2010-1 to 2012-1

^9^McMurdo Dry Valleys LTER - Beacon valley station; <https://www.mcmlter.org/> **|** 2010-1 to 2012-12

^(a)^Rainfall data were taken from 2017-2018 due to abnormal rainfall events after sample collection in 2016

^(b)^University Valley receives no liquid rainfall, all moisture is from humidity and meltwater

**Table S2: Weather data from sites.** Average, minimum and maximum weather data for a two-year period encompassing the sampling date of each site. Data sources and special notes are listed below the table.

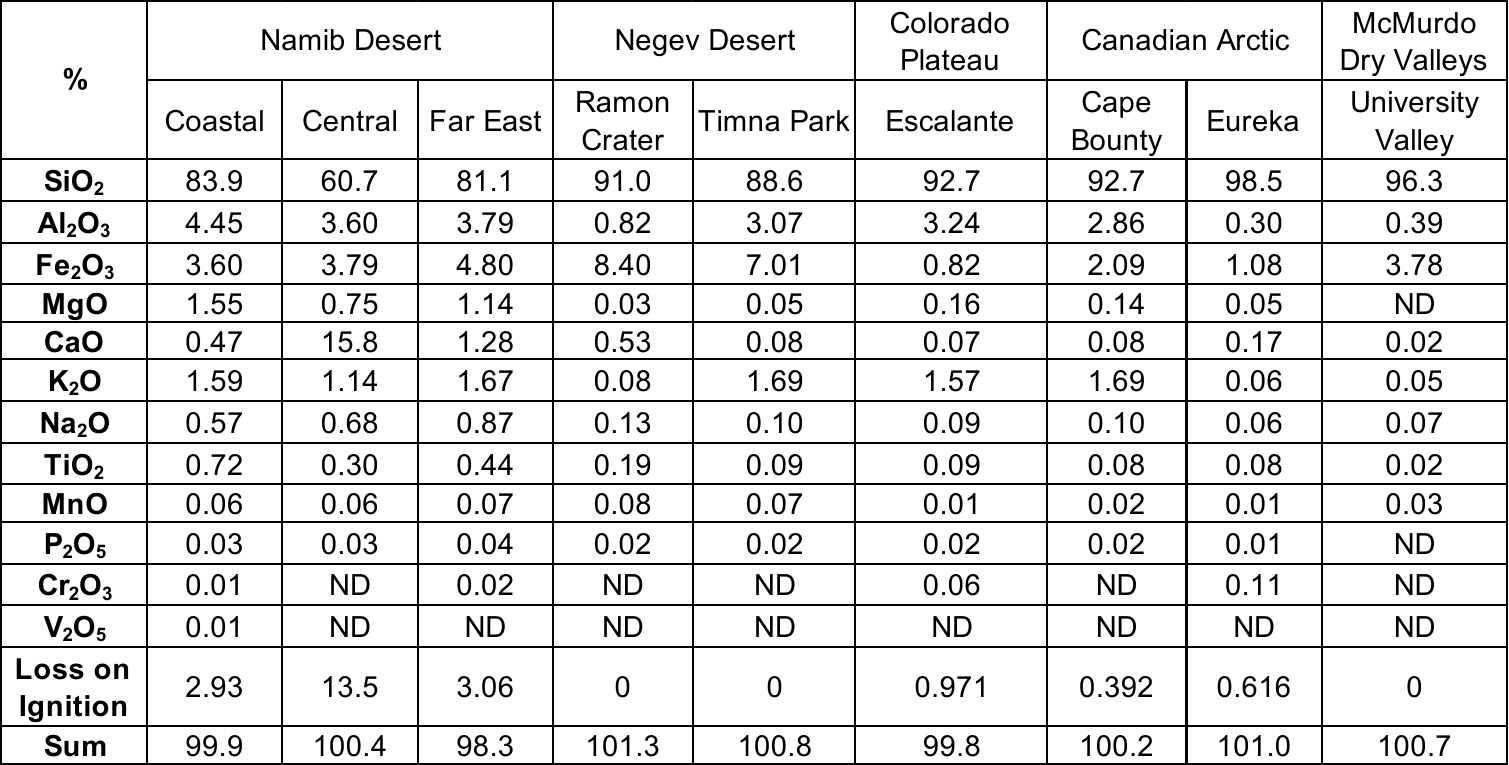

**Table S3: Chemical composition of sandstones.** Sandstone chemical composition was measured with X-ray fluorescence mass spectrometry. Loss on Ignition represents the percent of sandstone that was made of volatile substances. ND = below detection limit.

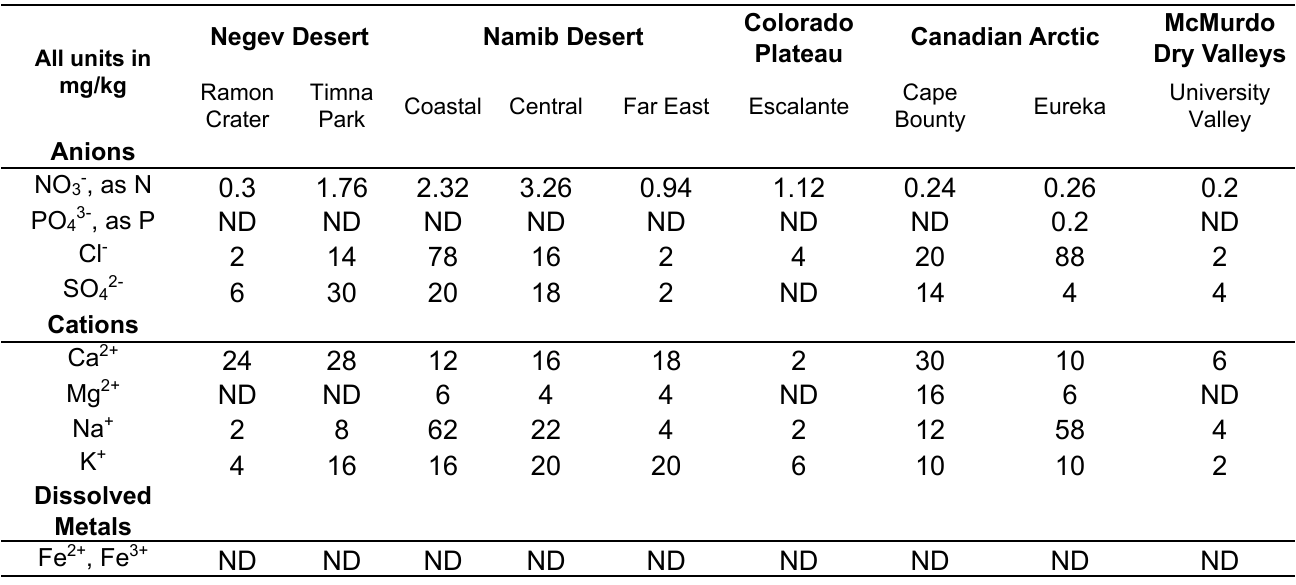

**Table S4: Water-soluble ions from sandstones.** Water soluble ions were extracted from crushed sandstone and measured with ion chromatography and inductively coupled plasma atomic emission spectroscopy. Units represent mg of ions per kg of rock. ND = below detection limit.

|  | **D50 Grain size (μm)** | **Percent water retention** |
| --- | --- | --- |
| Ramon Crater | 268 | 23.2 ± 1.7 |
| Timna Park | 530 | 14.9 ± 0.4 |
| Namib Coastal | 308 | 27.0 ± 1.3 |
| Namib Central | 233 | 11.0 ± 2.9 |
| Namib Far East | 172 | 30.3 ± 15.2 |
| Escalante | 150 | 11.8 ± 7.0 |
| Cape Bounty | 239 | 12.1 ± 2.1 |
| Eureka | 116 | 18.2 ± 1.1 |
| University Valley | 282 | 10.9 ± 0.7 |

**Table S5: Sandstone physical properties.** 50^th^ percentile diameter (D50) of grain size distribution and percent water retention measured with a water resaturation method.

|  | **Prokaryotic Diversity** | | **Eukaryotic Diversity** | |
| --- | --- | --- | --- | --- |
|  | OTUs | Shannon | ESVs | Shannon |
| Ramon Crater | 41 ± 5 | 4.57 ± 0.24 | - | - |
| Timna Park | 54 ± 7 | 5.11 ± 0.19 | - | - |
| Namib Coastal | 50 ± 8 | 4.57 ± 0.28 | - | - |
| Namib Central | 71 ± 4 | 5.61 ± 0.11 | - | - |
| Namib Far East | 92 ± 4 | 6.03 ± 0.07 | - | - |
| Escalante | 252 ± 14 | 7.51 ± 0.10 | 60 ± 13 | 3.28 ± 0.48 |
| Cape Bounty | 105 ± 17 | 6.03 ± 0.25 | 74 ± 14 | 4.46 ± 0.37 |
| Eureka | 125 ± 7 | 6.57 ± 0.09 | 29 ± 10 | 3.68 ± 0.27 |
| University Valley | 31 ± 5 | 4.37 ± 0.20 | 20 ± 2 | 3.26 ± 0.18 |

**Table S6: Alpha diversity by site.** Mean and standard deviation of observed OTUs/ESVs and Shannon index at each site, based on 16S rRNA gene sequences for prokaryotes and ITS sequences for eukaryotes.

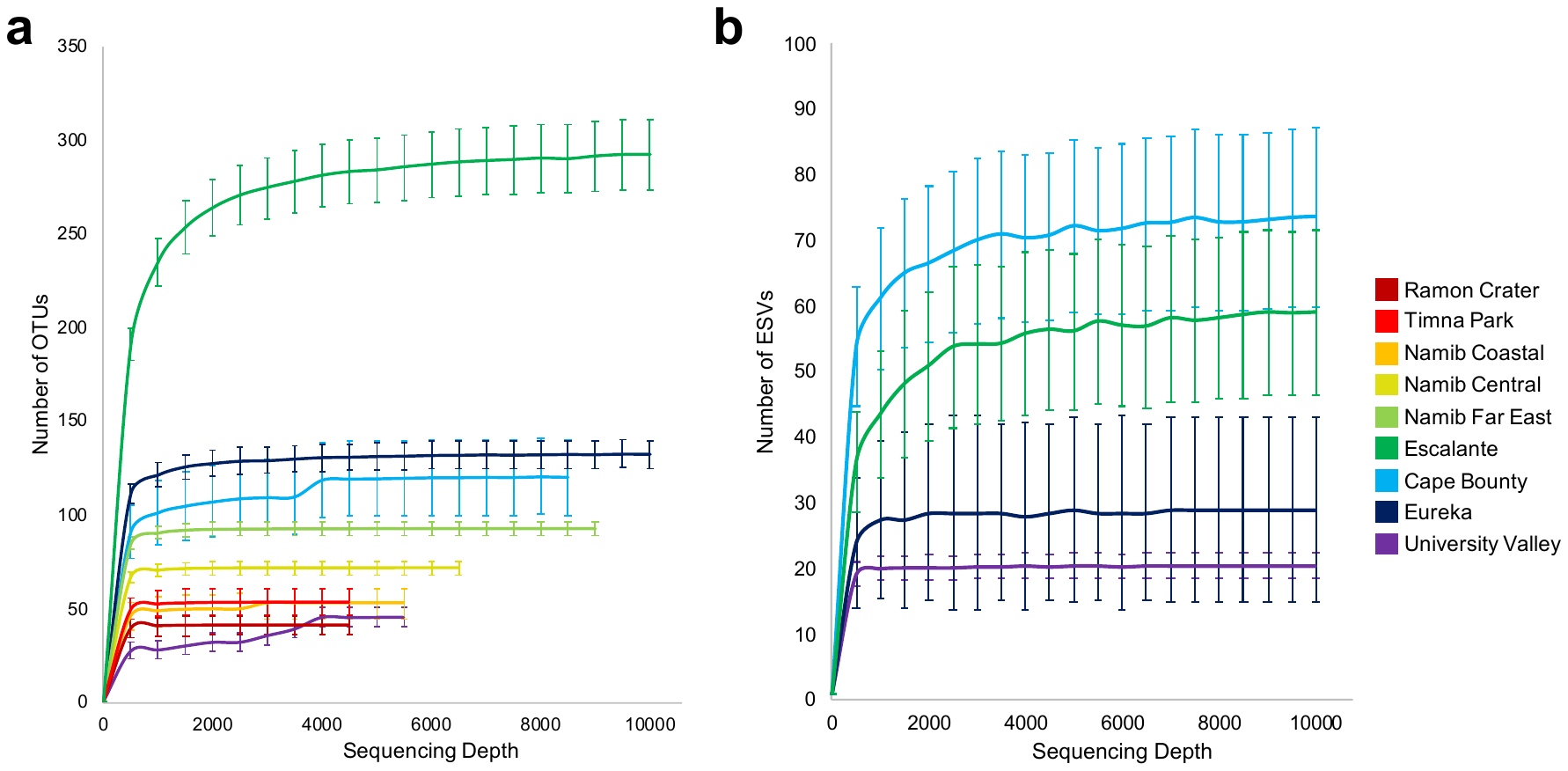

**Figure S1: Diversity rarefaction curves**. Rarefaction curves for **(a)** prokaryotic OTUs and **(b)** eukaryotic ESVs at 500-sequence sampling intervals, up to 10,000 sequences. These curves show that diversity approaches asymptote before maximum sequencing depth is reached

**Figure S2: Grain size distributions for sandstones**. Histograms showing grain size distributions for sandstones at each site.

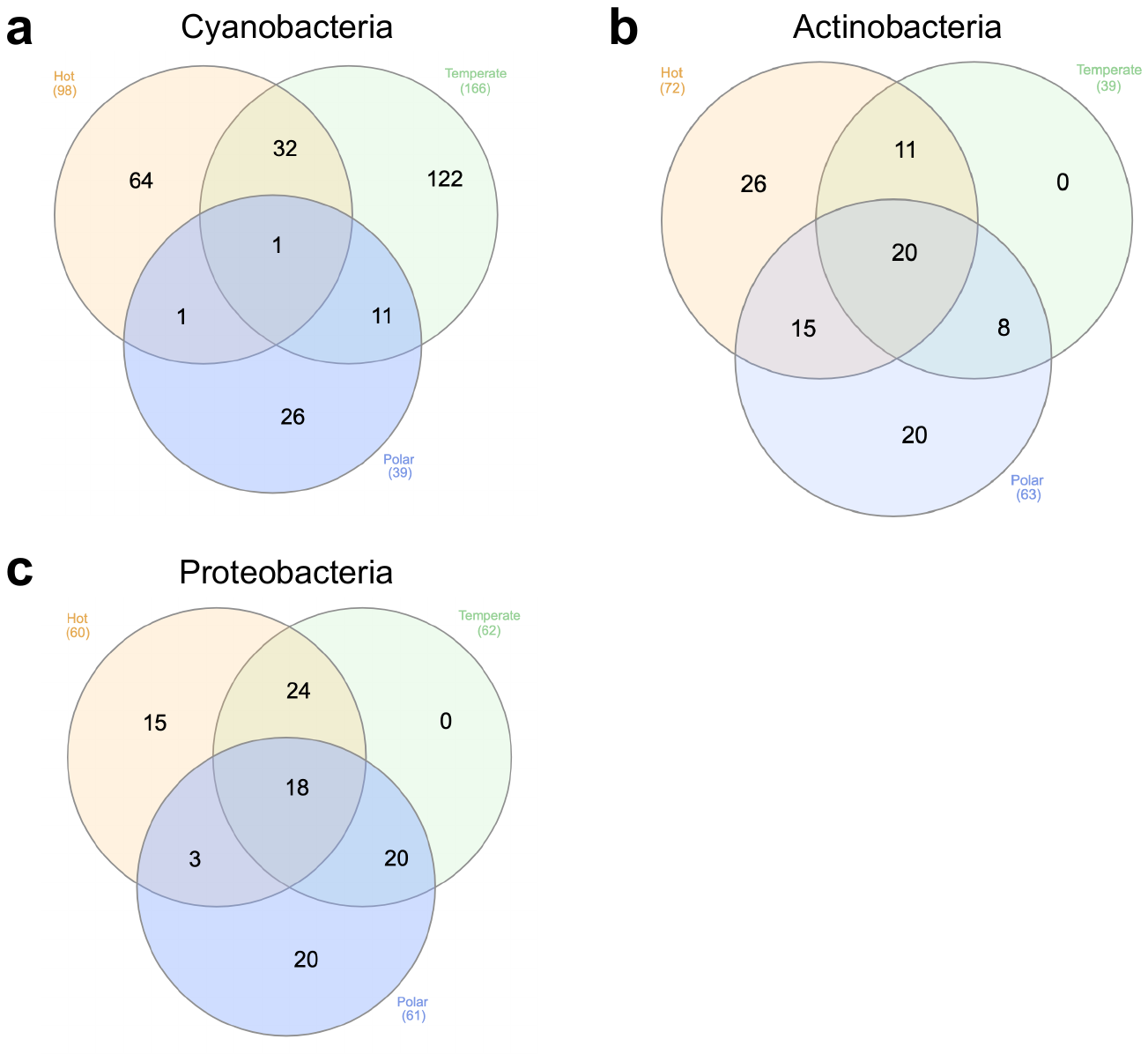

**Figure S3:** Venn diagram showing distribution of **(a)** all Cyanobacteria OTUs, **(b)** 100 most abundant Actinobacteria OTUs, and **(c)** 100 most abundant Proteobacteria OTUs between the three climate regimes (hot, polar, and temperate).

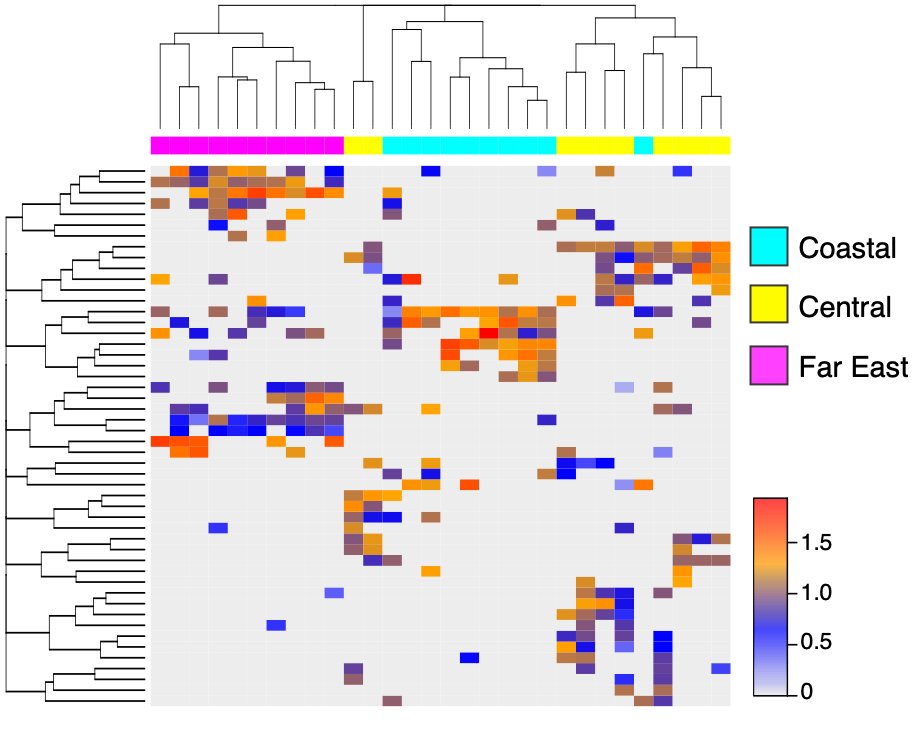

**Figure S4:** **Heatmap of** **Namib Desert Actinobacteria**. Actinobacteria OTUs from three sites in the Namib Desert. Rows correspond to OTUs and columns correspond to individual samples. Color scale represents log-normalized relative abundances.
